# Universal constraints on protein evolution in the long-term evolution experiment with *Escherichia coli*

**DOI:** 10.1101/2020.11.23.394791

**Authors:** Rohan Maddamsetti

## Abstract

Although it is well known that abundant proteins evolve slowly across the tree of life, there is little consensus for why this is true. Here, I report that abundant proteins evolve slowly in the hypermutator populations of Lenski’s long-term evolution experiment with *Escherichia coli* (LTEE). Specifically, the density of all observed mutations per gene, as measured in metagenomic time series covering 60,000 generations of the LTEE, significantly anti-correlates with mRNA abundance, protein abundance, and degree of protein-protein interaction. The same pattern holds for nonsynonymous mutation density. However, synonymous mutation density, measured across the LTEE hypermutator populations, positively correlates with protein abundance. These results show that universal constraints on protein evolution are visible in data spanning three decades of experimental evolution. Therefore, it should be possible to design experiments to answer why abundant proteins evolve slowly.

**Significance Statement:** A universal evolutionary pattern is that highly abundant and highly interacting proteins evolve slowly. This pattern was discovered in analyses that cover millions of years’ worth of sequence variation, so it is not clear how long it takes (decades, centuries, millennia) for such patterns to emerge. Here, I report that this universal evolutionary pattern emerges in metagenomic data that cover just 30 years of experimental evolution.

## Introduction

One consequence of the high complexity and intricate functional organization of organisms is that most mutations are deleterious. Natural selection resists the loss of function and fitness caused by mutation accumulation over time (Leiby and Marx 2014; LaBar and Adami 2017; Grant, et al. 2020). This process, called purifying selection, maintains the complexity and functional integrity of evolved organisms.

Despite its importance, purifying selection has been little studied in experimental systems (Alvarez-Ponce, et al. 2016), in contrast to adaptive evolution (Barrick and Lenski 2013). In two recent papers, my colleagues and I reported evidence for purifying selection in metagenomic time series of Lenski’s long-term evolution experiment with *Escherichia coli*, often called the LTEE for short (Lenski, et al. 1991; Good, et al. 2017). We considered the molecular evolution of the six hypermutator LTEE populations, which have elevated mutation rates due to evolved defects in DNA repair (Tenaillon, et al. 2016; Maddamsetti and Grant 2020a). These populations continue to increase in fitness due to adaptive evolution, even though genome evolution in these populations largely reflects the accumulation of nearly neutral mutations (Couce, et al. 2017). In Grant et al. (2020), we reported evidence for purifying selection on aerobic-specific and anaerobic-specific genes in *E. coli.* In Maddamsetti and Grant (2020b), we then reported evidence for purifying selection on genes that were found to be essential in the ancestral LTEE strain, REL606, in a transposon mutagenesis screen (Couce, et al. 2017).

Here, I report evidence that purifying selection in the LTEE reflects a universal constraint on protein evolution found across the tree of life, namely that highly abundant and highly interacting proteins evolve slowly (Fraser, et al. 2002; Hahn, et al. 2004; Drummond, et al. 2005; Hahn and Kern 2005; Drummond and Wilke 2008; Alvarez-Ponce, et al. 2017). Despite the universality and simplicity of this pattern of purifying selection, its proximate causes continue to be debated (Plata, et al. 2010; Plata and Vitkup 2018; Razban 2019; Usmanova, et al. 2020). A number of compelling hypotheses have been proposed, but consensus has not been reached. The findings reported here will not settle this debate. Nonetheless, an important consequence of my findings that is that it may be possible to resolve the causes of this universal pattern by experimental means.

## Results

### Rationale and study design

This study takes a novel approach to study the anti-correlation between protein abundance and evolutionary rates (Pál, et al. 2001; Drummond, et al. 2005; Drummond, et al. 2006; Drummond and Wilke 2008; Lobkovsky, et al. 2010; Yang, et al. 2010; Wylie and Shakhnovich 2011; Serohijos, et al. 2012; Serohijos and Shakhnovich 2014). In this section, I present the logical structure of the hypotheses and predictions under consideration, and explain the methods that I use (Figure 1).

**Figure 1.**
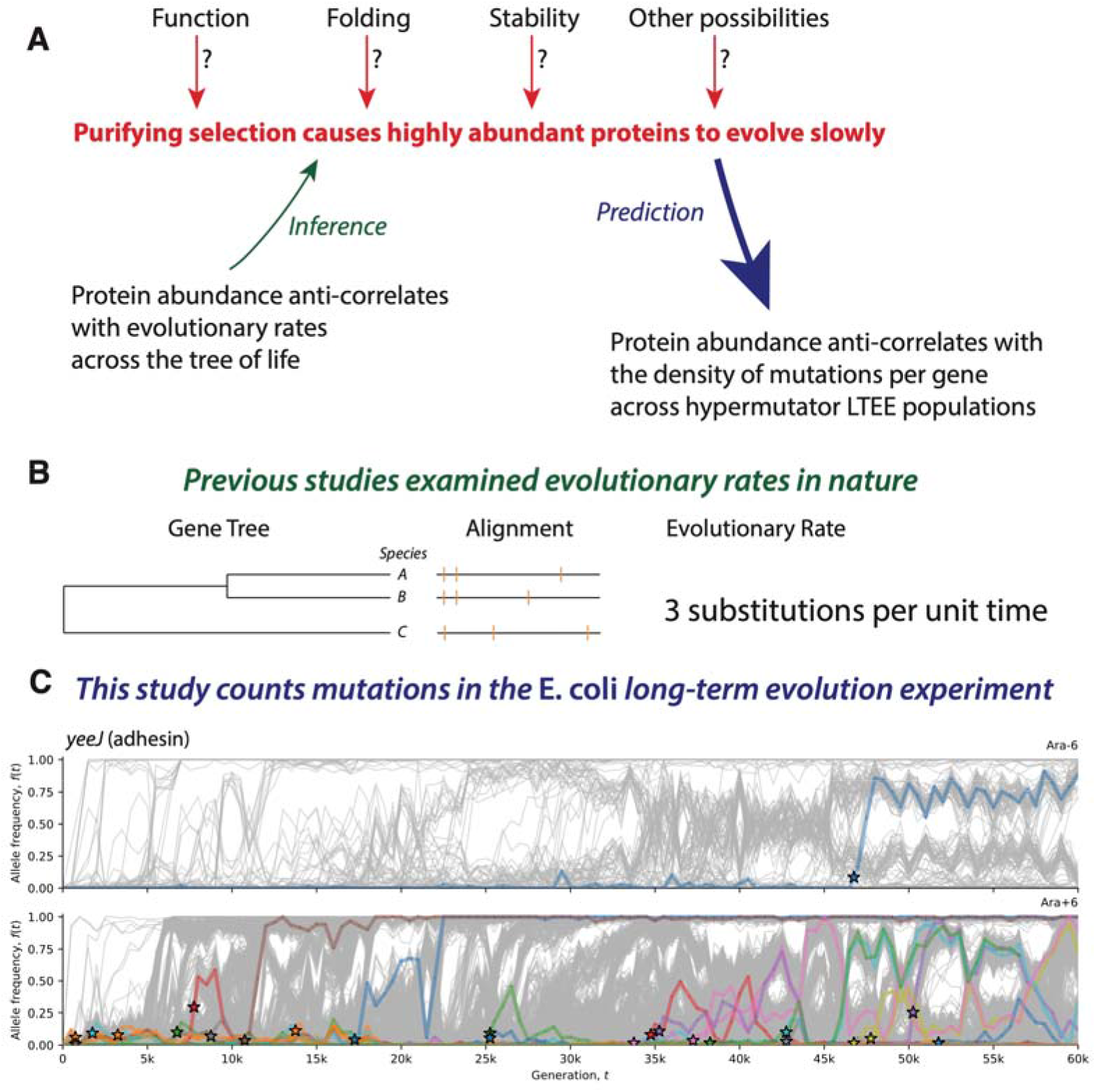
Study design. A) Many studies have reported that highly abundant proteins evolve slowly. If this fact is caused by purifying selection, then mutations in highly abundant proteins, should be more deleterious than mutations in less abundant proteins, on average. This logic leads to the prediction that highly abundant proteins should have fewer observed mutations than less abundant proteins across the hypermutator populations of the LTEE, taking gene length into account. B) Previous studies inferred evolutionary rates using DNA and protein sequence comparisons across species. C) This study sums all observed mutations per gene in metagenomic time series of the long-term evolution experiment with *Escherichia coli* (LTEE), considering nonmutator and hypermutator populations separately. This approach increases statistical power over a rate-based approach, and is neither affected by clonal interference nor frequencydependent selection. To give a concrete example, the top panel in (C) shows the number of observed mutations (stars) in the adhesin gene *yeeJ* in the nonmutator population Ara-6 over 60,000 generations. The bottom panel in (C) shows the number of observed mutations (stars) in *yeeJ* in the hypermutator population Ara+6 over the same period. For comparison across genes, the number of observed mutations is normalized by gene length.

I assume that the mutation rates in the hypermutator LTEE populations are high enough that the vast majority of observed mutations are nearly-neutral hitchhikers, whose dynamics are driven by a relatively small number of highly beneficial mutations (Barrick and Lenski 2009; Levy, et al. 2015; Maddamsetti, Lenski, et al. 2015; Tenaillon, et al. 2016; Couce, et al. 2017; Good, et al. 2017; Ba, et al. 2019; Maddamsetti and Grant 2020a). This allows us to infer information about mutation rates and biases (Couce, et al. 2017; Maddamsetti and Grant 2020a) even under environmental and population-genetic conditions that favor strong positive selection. It follows that the mutations observed across the nonmutator and hypermutator LTEE populations, to a large extent, reflect different parts of the distribution of mutation fitness effects (DFE) per gene.

With this assumption in hand, I start from the hypothesis that purifying selection causes abundant proteins to evolve slowly. This means that the DFE for abundant proteins should contain more deleterious mutations than the DFE for less abundant proteins, all else being equal. It follows that highly abundant proteins should have fewer observed mutations in the hypermutator LTEE populations, because it is unlikely that highly deleterious mutations will reach observable allele frequencies in the LTEE, given the population-genetic conditions of the LTEE (Good et al. 2017). This is the logical basis for using the hypermutator LTEE populations to test for purifying selection on abundant proteins.

The key technical trick is that we do not need to calculate evolutionary rates for the LTEE—in fact, we can completely ignore the phylogenetic structure of each population. Instead, we only need to count the number of observed mutations per gene across all hypermutator populations, and normalize by gene length (Figure 1). An additional benefit of this approach, is that the effects of clonal interference and frequency-dependent selection (Maddamsetti, Lenski, et al. 2015; Good, et al. 2017) can be ignored, because these phenomena do not affect the density of mutations that are *ever* observed in the LTEE. By contrast, clonal interference and frequencydependent selection may have significant effects on evolutionary rates (Lang, et al. 2013; Serohijos and Shakhnovich 2014; Maddamsetti, Lenski, et al. 2015; Good, et al. 2017).

The great advantage of the LTEE, and other evolution experiments with microbes, is the “fossil record” of frozen samples that can be revived for comparison with later samples. The vast majority of mutations in the LTEE lie off the line of descent, but are still accessible from sequencing those frozen population samples (Good et al. 2017). By contrast, analyses of natural sequence data are largely restricted to extant within-population polymorphism and between-species fixations. The use of mutations off the line of descent in the LTEE, along with its multidecade duration, provides sufficient (and ever increasing) statistical power to discern patterns of purifying selection, such as the one discussed in this work.

### Correlations between mRNA and protein abundance and mutation density per gene in LTEE populations

I compared the density of observed mutations in the LTEE (Good, et al. 2017) to mRNA and protein abundance data for the LTEE ancestral strain, REL606, grown in DM500 media (Caglar, et al. 2017). These comparisons are shown in Figure 2; note that throughout this section, all significant Spearman correlation coefficients and associated *p*-values are labeled on the figures. In the hypermutator LTEE populations, mRNA abundance during exponential growth significantly anticorrelates with mutation density, while protein abundance, at all time points, significantly anticorrelates with mutation density. The same anti-correlation holds, for all time points, when only nonsynonymous (i.e., missense and nonsense) mutations are considered (Figure 3). The significance of these anti-correlations increases when genes with no observed mutations in the metagenomic data are excluded (Supplementary Figure S1 for all mutation types; Supplementary Figure S2 for nonsynonymous mutations). By contrast, the density of synonymous mutations across the hypermutator populations shows a significant positive correlation with mRNA and protein abundance for REL606 in DM500 media, across all phases of growth (Figure 4). When genes with no mutations in the metagenomic data are excluded, significant positive correlations remain between synonymous mutation density and mRNA and protein abundance, although to a lesser degree (Supplementary Figure S3). In the nonmutator LTEE populations, both mRNA and protein abundance for REL606 grown in DM500 show significant positive correlations with the density of observed mutations (Supplementary Figure S4).

**Figure 2.**
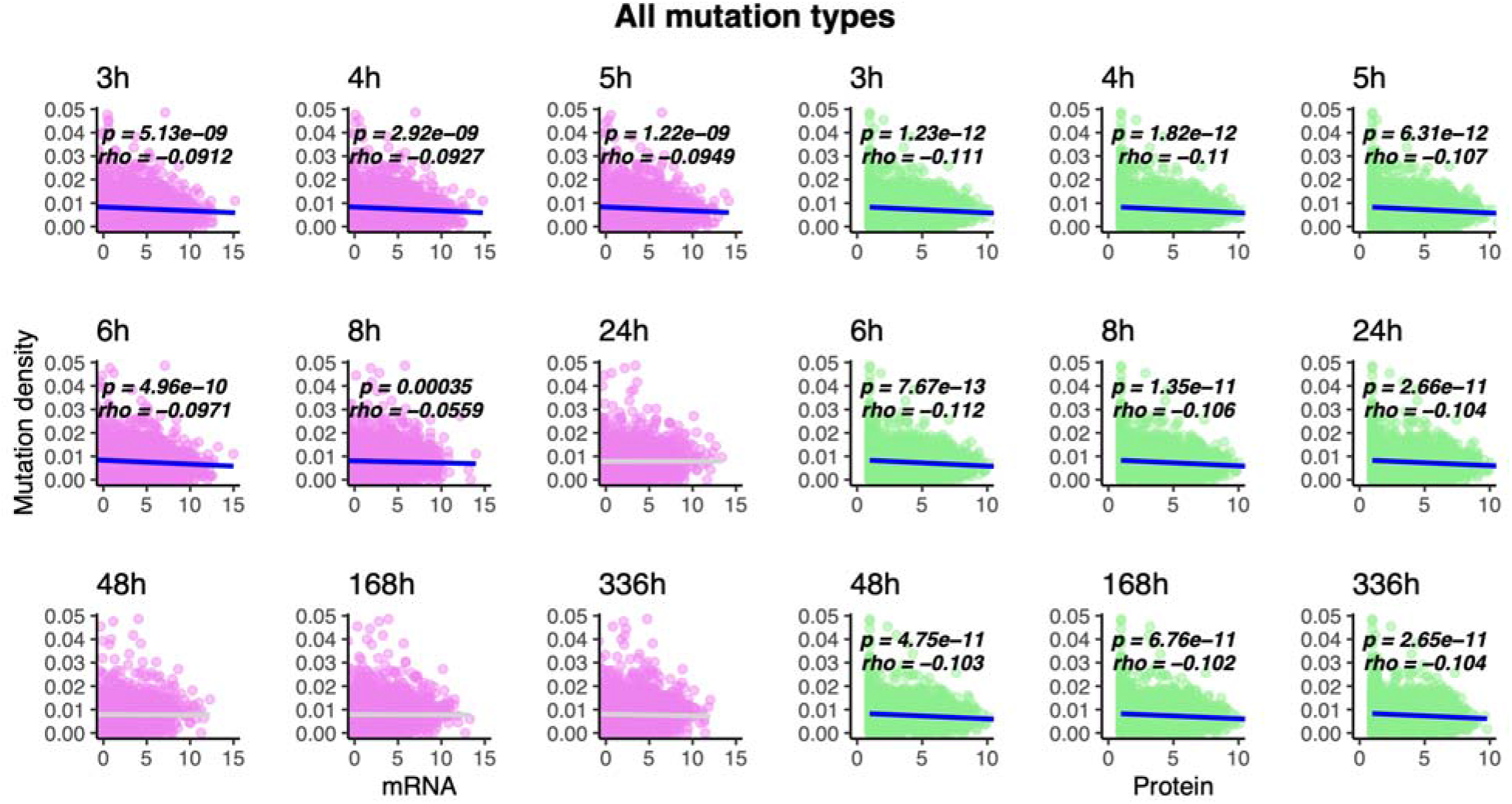
The density of observed mutations per gene across all hypermutator LTEE populations anti-correlates with mRNA abundance in exponential growth phase, and anticorrelates with protein abundance at all time points. RNA and protein abundance were measured for the ancestral LTEE clone REL606, grown in DM500 media (Caglar et al. 2017). Each point represents a protein-coding gene in the genome of the ancestral LTEE clone, *E. coli* B strain REL606. The abundance of mRNA or protein expressed per gene is shown on the x-axis of each plot. The density of observed mutations per gene is shown on the y-axis of each plot. Comparisons to mRNA abundance are shown in purple, while comparisons to protein abundance are shown in green. Statistically significant correlations are shown in blue, while non-significant correlations are shown in light gray. Spearman correlation coefficients (*rho*) and associated *p-* values are shown on each panel.

**Figure 3.**
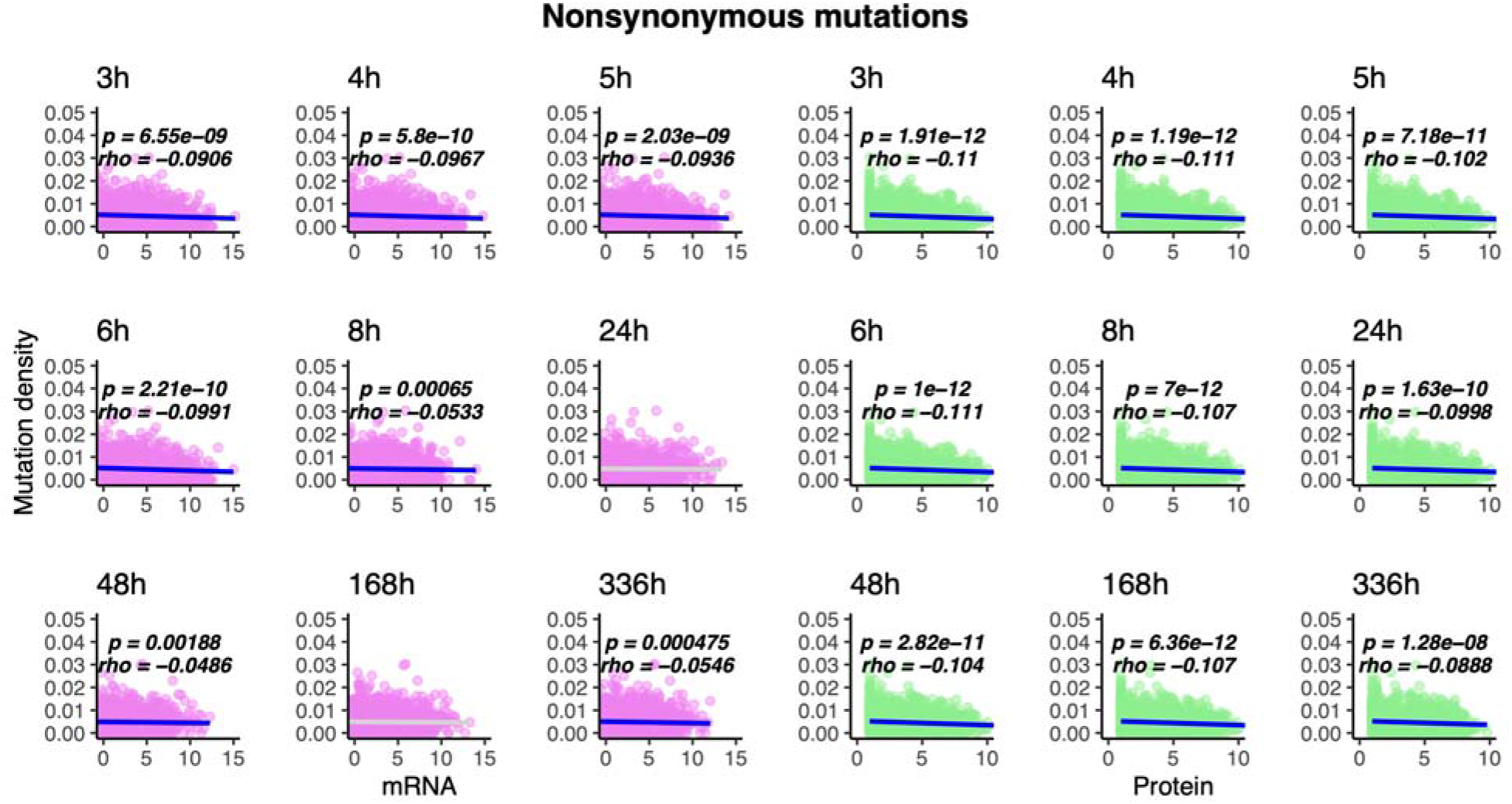
The density of observed nonsynonymous mutations per gene across all hypermutator LTEE populations anti-correlates with mRNA abundance in exponential growth phase, and anti-correlates with protein abundance at all time points. See Figure 2 legend for further details.

**Figure 4.**
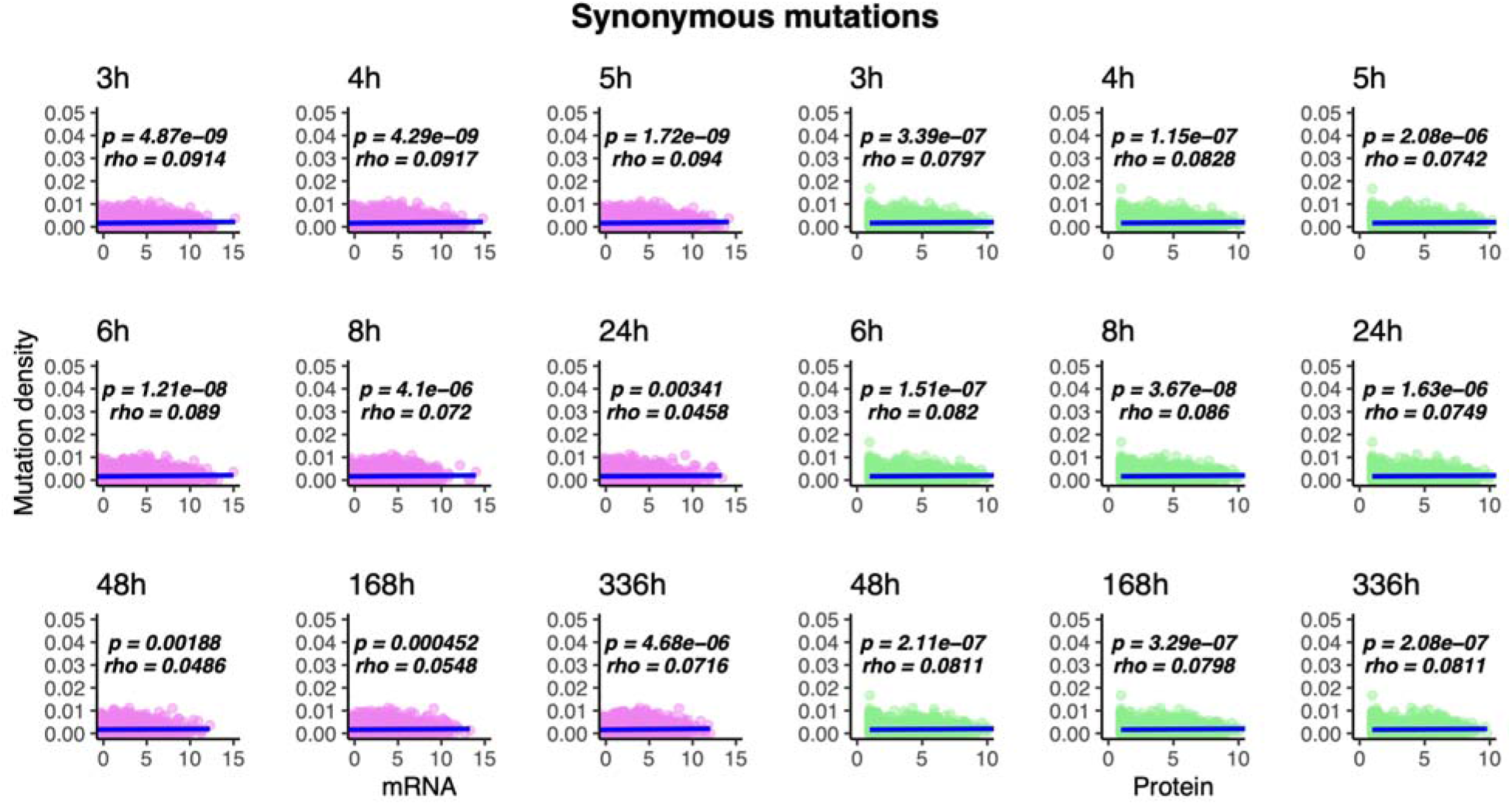
The density of observed synonymous mutations per gene across all hypermutator LTEE populations positively correlates with mRNA and protein abundance at all time points. See Figure 2 legend for further details.

I also asked whether the strength of the Spearman correlations between protein abundance and mutation density in the hypermutator populations increased over the course of the LTEE (Figure 5). In analyses of natural sequence variation, it is understood that the strength of anti-correlation between protein evolutionary rates and protein abundance increases with divergence time among the taxa under consideration (Serohijos, et al. 2012). Based on protein biophysics, Serohijos et al. (2012) additionally predicted that the strength of the anti-correlation between evolutionary rates and protein abundance would increase, but at declining rates over time. Even though the differences in measurements, units, and timescales make direct comparisons to those theoretical predictions impossible, it is striking that a similar functional form of the relationship between time and the strength of the rate-abundance anti-correlation occurs with the mutations observed across the LTEE hypermutator populations (Figure 5A and B). By contrast, the positive Spearman correlation coefficient between synonymous mutation density and protein abundance remains steady at ~0.075 for at least 40,000 generations, ranging from the 20,000-generation mark through 60,000 generations (Figure 5C).

**Figure 5.**
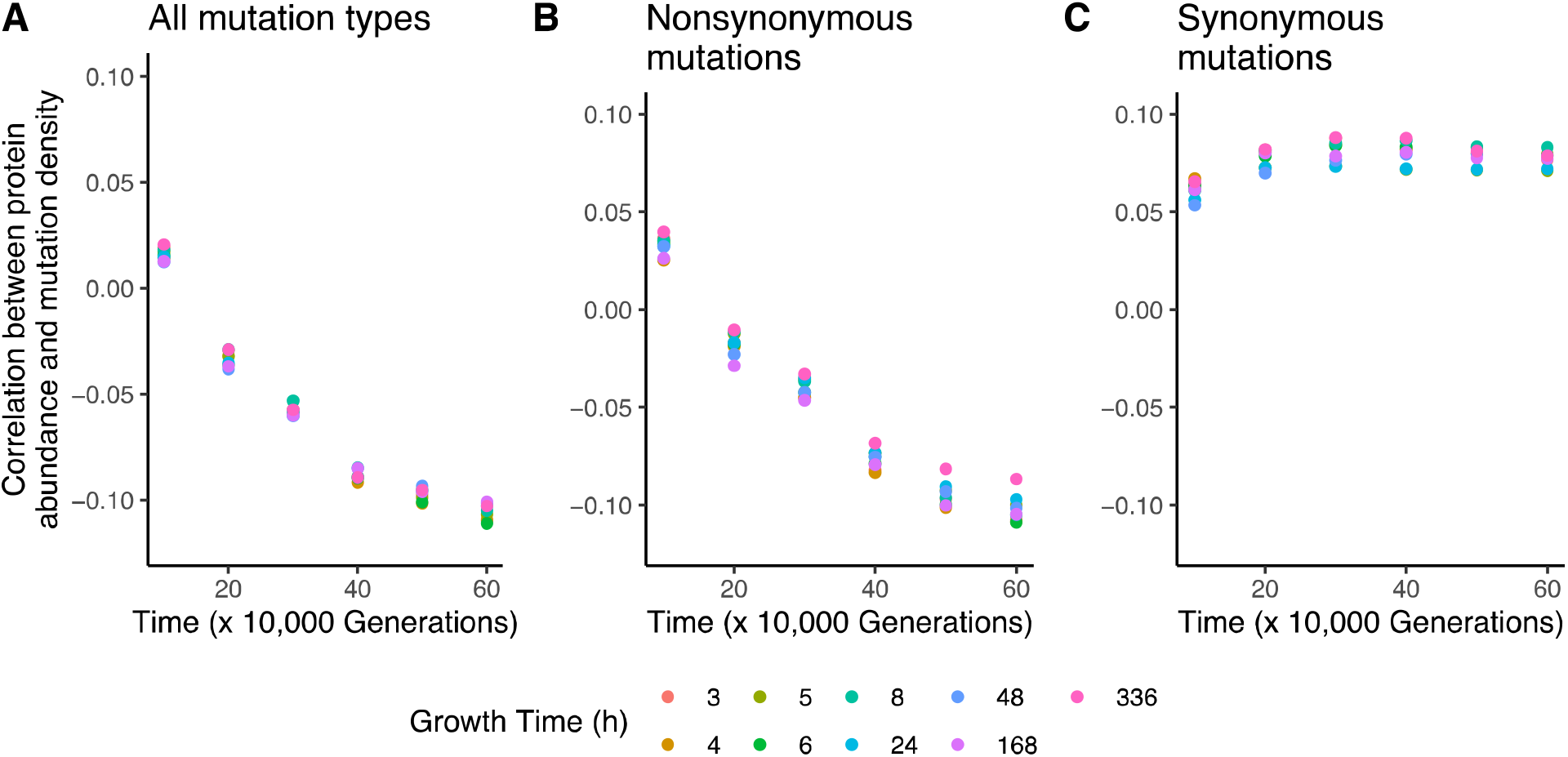
Correlations between protein abundance in REL606 and mutation density are consistent over time. Points represent Spearman correlation coefficients, calculated using the cumulative number of mutations observed by each 10,000-generation mark in the metagenomic time series for the LTEE hypermutator populations. Colors indicate the growth time at which protein abundance was sampled for REL606; the growth times correspond to the separate panels in Figures 2, 3, and 4. A) Correlations between protein abundance in REL606 and mutation density across all hypermutator LTEE populations. B) Correlations between protein abundance in REL606 and nonsynonymous mutation density across all hypermutator LTEE populations. C) Correlations between protein abundance in REL606 and synonymous mutation density across all hypermutator LTEE populations.

A limitation of these analyses is that these RNA and protein abundance data come from the ancestral LTEE clone, REL606, and so these patterns may not hold for evolved strains. To address this limitation, I examined RNA abundance data for 11 50,000 generation LTEE clones, grown to exponential phase in DM4000 media (Favate, et al. 2021). In every single case, the density of observed mutations per gene, measured across all hypermutator populations, significantly anti-correlates with mRNA abundance (Supplementary Figure S5). In addition, a significant anti-correlation is seen for nonsynonymous mutations in all 11 clones (Supplementary Figure S6), while a positive correlation is seen for synonymous mutations, again for all 11 evolved clones (Supplementary Figure S7). The density of observed mutations per gene in the nonmutator populations significantly correlates with mRNA abundance in 7 out of 11 clones (Supplementary Figure S8).

As an additional check for the robustness of these correlations, I compared the density of observed mutations per gene in the LTEE to protein abundance data in the ProteomeVis database (Razban, et al. 2018). Although these data only cover 664 out of 4,205 genes analyzed in the LTEE metagenomic data, they still reveal significant anti-correlations between mutation density per gene in the hypermutator populations and protein abundance, when all mutations and nonsynonymous mutations are analyzed (Supplementary Figure S9). Corresponding results for synonymous mutations in the hypermutator LTEE populations, and for all mutation types in the nonmutator LTEE populations, are not statistically significant.

### Highly interacting proteins evolve slowly in hypermutator populations

Another universal pattern is that highly interacting proteins evolve more slowly than those with fewer interaction partners (Fraser, et al. 2002; Hahn, et al. 2004; Hahn and Kern 2005; Alvarez-Ponce, et al. 2017). I hypothesized that highly interacting proteins would be under strong selection in the LTEE, based on those reports, as well as previous results showing that the *E. coli* core genome is under positive selection in the LTEE (Maddamsetti, et al. 2017), and that global regulators of gene expression show evidence of strong positive selection in both nonmutator and hypermutator LTEE populations (Maddamsetti and Grant 2020b). In particular, I hypothesized that highly interacting proteins should evolve rapidly in the nonmutator LTEE populations due to positive selection, but should evolve slowly in the hypermutator populations during to purifying selection.

I compared the number of protein-protein interactions (PPI) to the density of observed mutations across LTEE populations for every protein-coding gene in the *E. coli* genome, using three curated datasets of protein-protein interactions in *E. coli* (Razban, et al. 2018; Cong, et al. 2019; Zitnik, et al. 2019), which I refer to as the Cong dataset, the Zitnik dataset, and the Razban dataset. These comparisons are shown in Figure 6 and Supplementary Figure 10. I find significant negative correlations between mutation density and PPI degree in the hypermutators (Spearman’s *rho* = −0.056,*p* = 0.00037 for Cong dataset; Spearman’s *rho* = −0.11,*p* < 10^−11^ for Zitnik dataset; Spearman’s *rho* = −0.068,*p* < 10^−4^ for Razban dataset). However, the weak positive correlations between mutation density and protein-protein interaction degree (PPI degree) in the nonmutators are not significant (Supplementary Figure 10).

**Figure 6.**
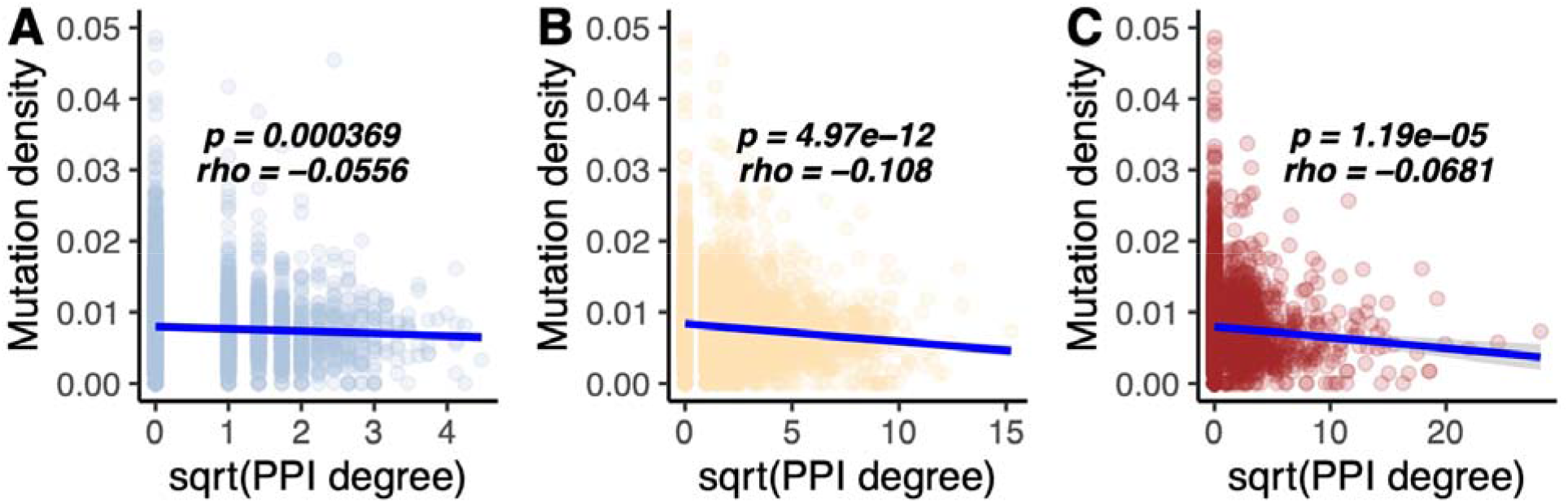
The density of observed mutations per gene across all hypermutator LTEE populations negatively correlates with PPI degree. Comparisons to the PPI data from Cong et al. (2019) are shown in light blue, comparisons to the PPI data from Zitnik et al. (2019) are shown in orange, and comparisons to the PPI data in the ProteomeVis database (Razban et al. 2018) are shown in red. Significant Spearman correlations are shown in blue. For improved visual dispersion, PPI degree is square-root transformed; the Spearman correlation is unaffected by this monotonic data transformation. A) Proteins with more interactions in the Cong et al. (2019) dataset tend to evolve more slowly than those with fewer interactions in the hypermutator LTEE populations. B) Proteins with more interactions in the Zitnik et al. (2019) dataset tend to evolve more slowly than those with fewer interactions in the hypermutator LTEE populations. C) Proteins with more interactions in the ProteomeVis *E. coli* PPI dataset (Razban et al. 2018) tend to evolve more slowly than those with fewer interactions in the hypermutator LTEE populations.

## Discussion

I show that a number of well-known but poorly understood correlations between mRNA abundance, protein abundance, PPI degree, and evolutionary rates across the tree of life are also found in the hypermutator populations of the LTEE. In some cases, I find significant anticorrelation between mutation densities and mRNA abundance in exponential phase, but not during stationary phase. The simplest explanation for this finding is that mRNAs decay more rapidly than the proteins they encode. Protein abundance consistently shows a consistent negative correlation with the density of all observed mutations (Figure 2) and with nonsynonymous mutation density across all time points (Figure 3).

It is widely believed that these correlations are driven by purifying selection on universal aspects of protein evolution (Drummond, et al. 2006; Drummond and Wilke 2009; Serohijos, et al. 2012; Serohijos and Shakhnovich 2014), and indeed, this is the most parsimonious explanation for why similar patterns are seen in the LTEE. An intriguing difference, however, is the positive correlation that I find between synonymous mutation density across LTEE hypermutator populations and protein abundance (Figure 4) — which contrasts with the anticorrelation between the rate of synonymous mutations and gene expression seen in nature (Drummond and Wilke 2008). In part, this may be explained by the differences in the distribution of synonymous mutations observed in the LTEE, and the distribution of synonymous diversity per gene in nature (Maddamsetti, Hatcher, et al. 2015; Maddamsetti and Grant 2020a), although the causes for this difference between natural variation and experiment is still a matter for hypothesis generation (Maddamsetti 2016), data collection, hypothesis testing, and debate.

An important limitation of these results is that the protein and mRNA abundance data for LTEE strains were collected in DM500 and DM4000 media (Caglar, et al. 2017; Favate, et al. 2021). These media contain much more than the 25 mg/L glucose in the DM25 media used in the LTEE. This represents a technical compromise due to the fact that researchers have not yet succeeded in isolating sufficient mRNA from exponential phase cultures in DM25 for RNA-Seq (Tanush Jagdish and Nkrumah Grant, personal communication). With this caveat in mind, my findings support the conclusion that highly abundant proteins evolve slowly in the hypermutator LTEE populations.

The causes for why highly abundant proteins evolve slowly may emerge from a number of different, and non-mutually exclusive phenomena, so many explanations have been proposed (Razban 2019). These include the protein misfolding avoidance hypothesis (Yang, et al. 2010), the protein misinteraction avoidance hypothesis (Levy, et al. 2012; Yang, et al. 2012), the mRNA folding hypothesis (Park, et al. 2013), purifying selection on protein function (Konaté, et al. 2019), folding stability (Serohijos, et al. 2012; Serohijos and Shakhnovich 2014), and others (Tartaglia, et al. 2007; Plata, et al. 2010; Kepp and Dasmeh 2014).

Differentiating among these possibilities is difficult, because it is challenging to study the *causes* of patterns that span millions of years of protein evolution. I do not draw conclusions about the causes of these correlations. Rather, my results show that evolution experiments are reasonable model systems to study the causes of evolutionary rate variation in proteins. A concrete approach would be to recode the genomes of hypermutator strains to modulate the anticipated action of purifying selection per protein, based on the predictions of a particular explanation, and then ask whether those predictions are borne out during experimental evolution. Breakthroughs that allow for the inexpensive recoding of whole bacterial genomes may be needed, but it is plausible that such experiments will be feasible in the future.

Many other experimental directions are possible. First, a better understanding of how chaperones and other molecular mechanisms of protein quality control affect evolutionary rates and fitness (Chen, et al. 2017; Alvarez-Ponce, et al. 2019; Samhita, et al. 2020) is needed. We also need to better understand purifying selection on synonymous mutations (Walsh, et al. 2020). Second, studies on how RNA transcription error rates (Li and Lynch 2020) and RNA folding errors affect evolutionary rates would be valuable. Indeed, mRNA accessibility seems to be an important predictor of protein abundance (Terai and Asai 2020). Third, it would be interesting to experimentally test the hypothesis that protein and RNA chaperones evolve under more and more stringent purifying selection during long-term experimental evolution, which follows from the premise that hypermutator LTEE populations are affected by a mutation load that affects protein folding and stability. Studies on the existence and relevance of phenomena like evolutionary capacitance caused by the contributions that protein-protein interactions make to folding stability (Dixit and Maslov 2013; Jarzab, et al. 2020; Mateus, et al. 2020), including cryptic genetic variation hidden by protein and RNA chaperones (Queitsch, et al. 2002; Bergman and Siegal 2003; Masel 2005, 2006, 2013; Trotter, et al. 2014; Geiler-Samerotte, et al. 2016; Zheng, et al. 2019) during experimental evolution, and the effects of such phenomena on rates of protein evolution may be especially valuable in this regard. Finally, it would be valuable to develop a better understanding of the temperature sensitivity of evolved LTEE populations (Mongold, et al. 1996, 1999; Leiby and Marx 2014), and to collect data on protein evolutionary rates in long-term experiments conducted at elevated temperatures (Bennett, et al. 1990; Tenaillon, et al. 2012). Much remains to be explored, in regard to how evolution experiments can deepen our understanding of purifying selection on molecular and cellular organization and function.

## Materials and Methods

Pre-processed LTEE metagenomic data was downloaded from: https://github.com/benjaminhgood/LTEE-metagenomic. Transcriptomic and proteomic data for REL606, grown in Davis minimal media with 500 mg/L glucose (DM500), was taken from the supplementary tables for Caglar et al. (2017). For robustness, I also analyzed the transcriptomic data for 11 50,000 generation LTEE clones grown in DM4000 media (Favate, et al. 2021) available at: https://github.com/shahlab/LTEE-gene-expression. I analyzed three different datasets of protein-protein interactions (PPI) in *E. coli.* First, I used the PPI network for *E. coli* K-12 MG1655 in the STRING database (Szklarczyk, et al. 2016) as curated by Zitnik et al. (2019). Second, I used the dataset of high confidence *E. coli* PPI interactions reported by Cong et al. (2019), which combines co-evolutionary information in large protein multiple sequence alignments with gold-standard protein complexes in *E. coli* reported in the Ecocyc and Protein Databank (PDB) databases (Berman, et al. 2000; Keseler, et al. 2013). PPI network statistics were calculated using the SNAP toolkit (Leskovec and Sosič 2016; Zitnik, et al. 2019). Third, additional data on *E. coli* PPI interactions and protein abundance was downloaded using the web interface to the ProteomeVis database (Razban, et al. 2018), available at http://proteomevis.chem.harvard.edu/. Associated metadata for ProteomeVis was downloaded from: https://github.com/rrazban/proteomevis/blob/master/make_database/proteomevis_inspect.csv.

All statistical analyses involve two-sided tests for Spearman correlation coefficients that are significantly different from zero, using the cor.test function in the R statistical programming language, version 4.0 (R Core Team 2020). Unless stated otherwise, all correlations include genes with no mutations (i.e., zeros are included).

## Data Availability Statement

The data and analysis codes underlying this article are available on the Dryad Digital Repository (DOI pending publication). Analysis codes are also available at: https://github.com/rohanmaddamsetti/LTEE-network-analysis.

## Acknowledgements

I thank Richard Lenski, Lingchong You, Jeffrey Barrick, Stephanie Spielman, Tanush Jagdish, and Nkrumah Grant for valuable comments, advice, and discussions. I thank Rostam Razban for help with the ProteomeVis database and John Favate and Premal Shah for access and discussions about their LTEE RNA-seq data. The LTEE that generated the bacteria used in this study is supported by a grant from the National Science Foundation (currently DEB-1951307) to Richard Lenski and Jeffrey Barrick.

## SUPPLEMENTARY MATERIAL

**Supplementary Figure S1.**
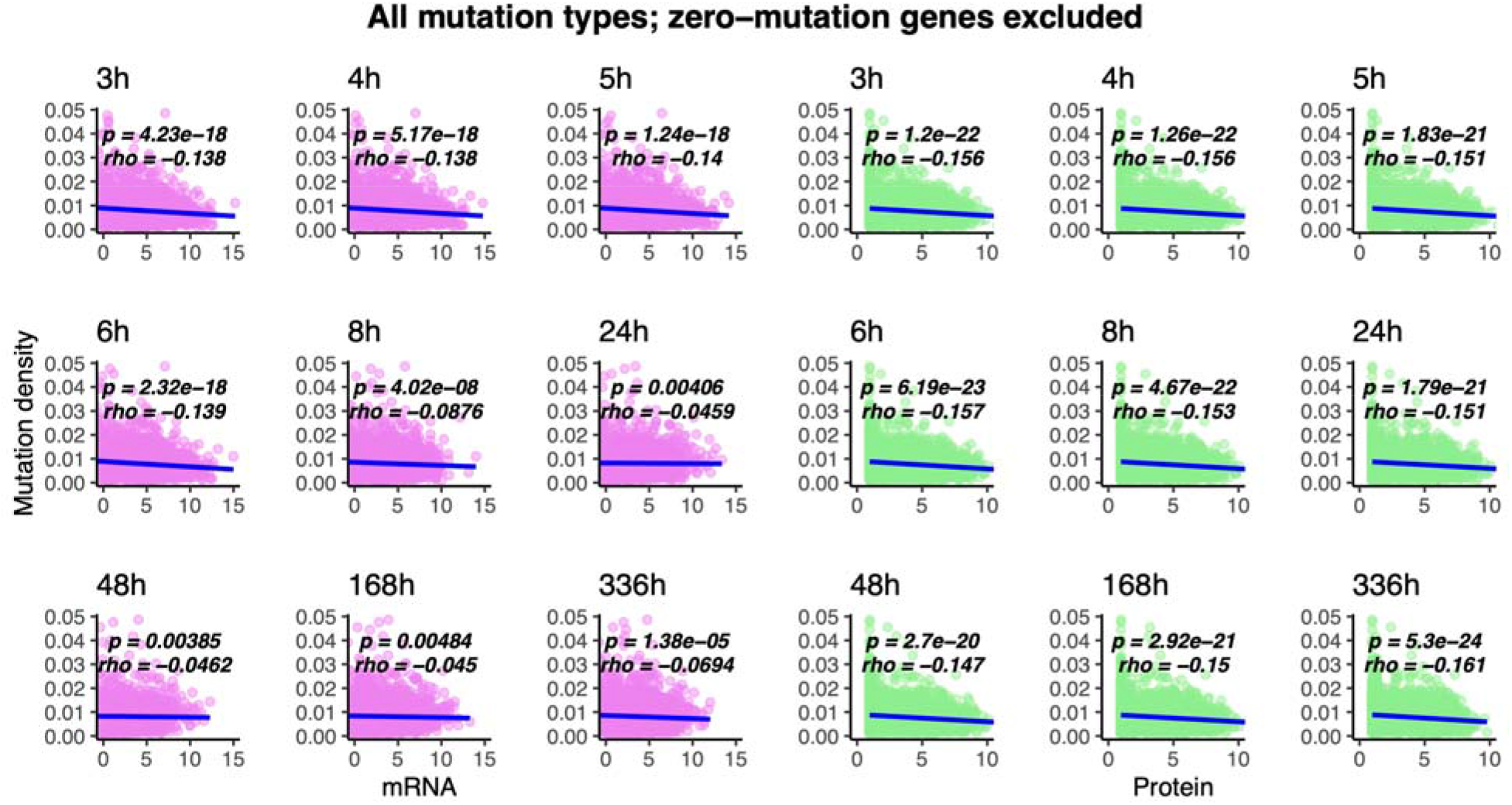
The density of observed mutations per gene across all hypermutator LTEE populations anti-correlates with mRNA and protein abundance at all time points, when genes with no mutations are excluded. RNA and protein abundance were measured for the ancestral LTEE clone REL606, grown in DM500 media (Caglar et al. 2017). Each point represents a protein-coding gene in the genome of the *E. coli* B strain REL606. The abundance of mRNA or protein expressed per gene is shown on the x-axis of each plot. The density of observed mutations per gene is shown on the y-axis of each plot. Comparisons to mRNA abundance are shown in purple, while comparisons to protein abundance are shown in green. Statistically significant linear regressions are shown in blue, while non-significant regressions are shown in light gray.

**Supplementary Figure S2.**
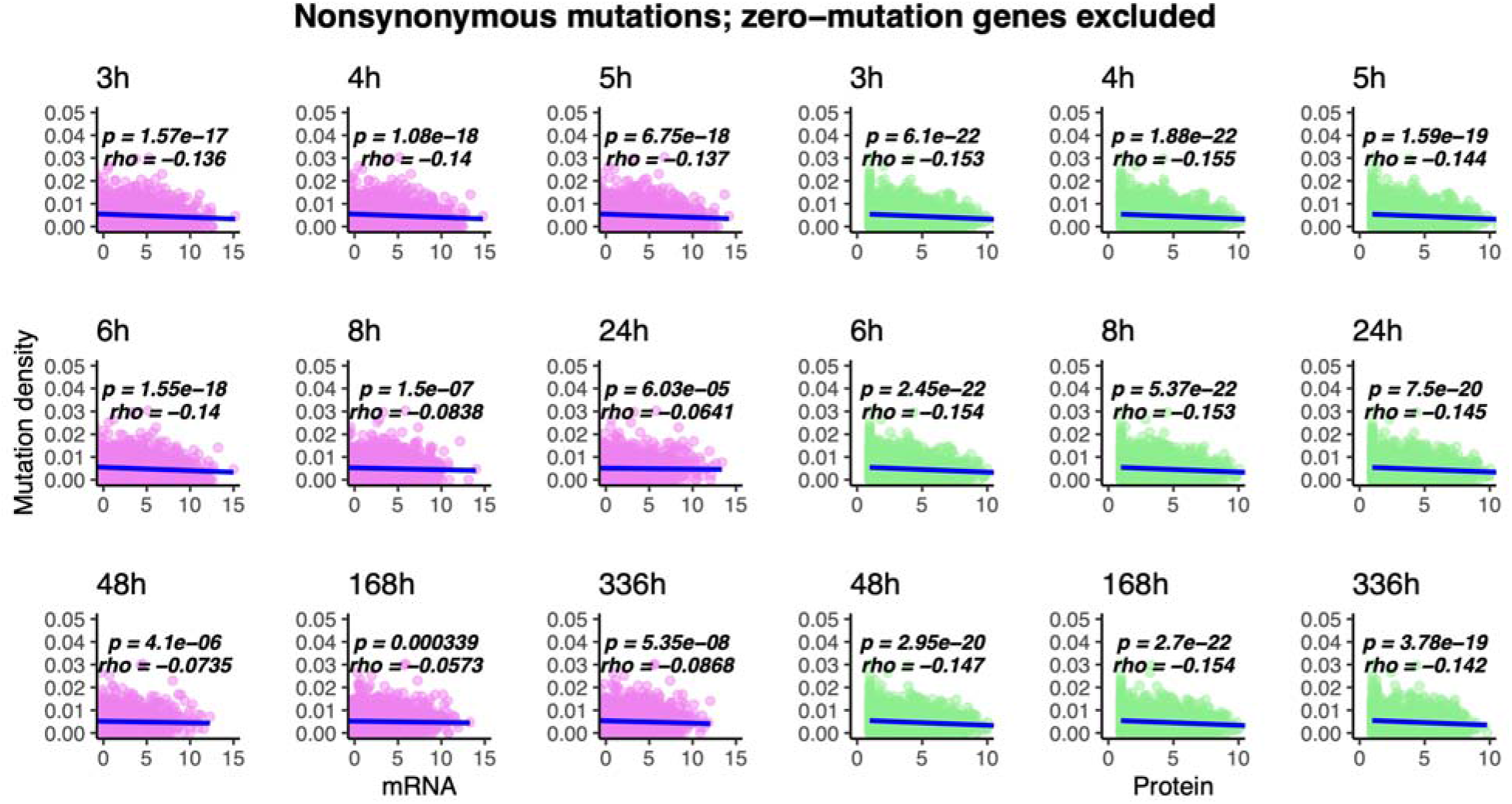
The density of observed nonsynonymous mutations per gene across all hypermutator LTEE populations anti-correlates with mRNA and protein abundance at all time points, when genes with no mutations are excluded. See legend to Supplementary Figure S1 for further details.

**Supplementary Figure S3.**
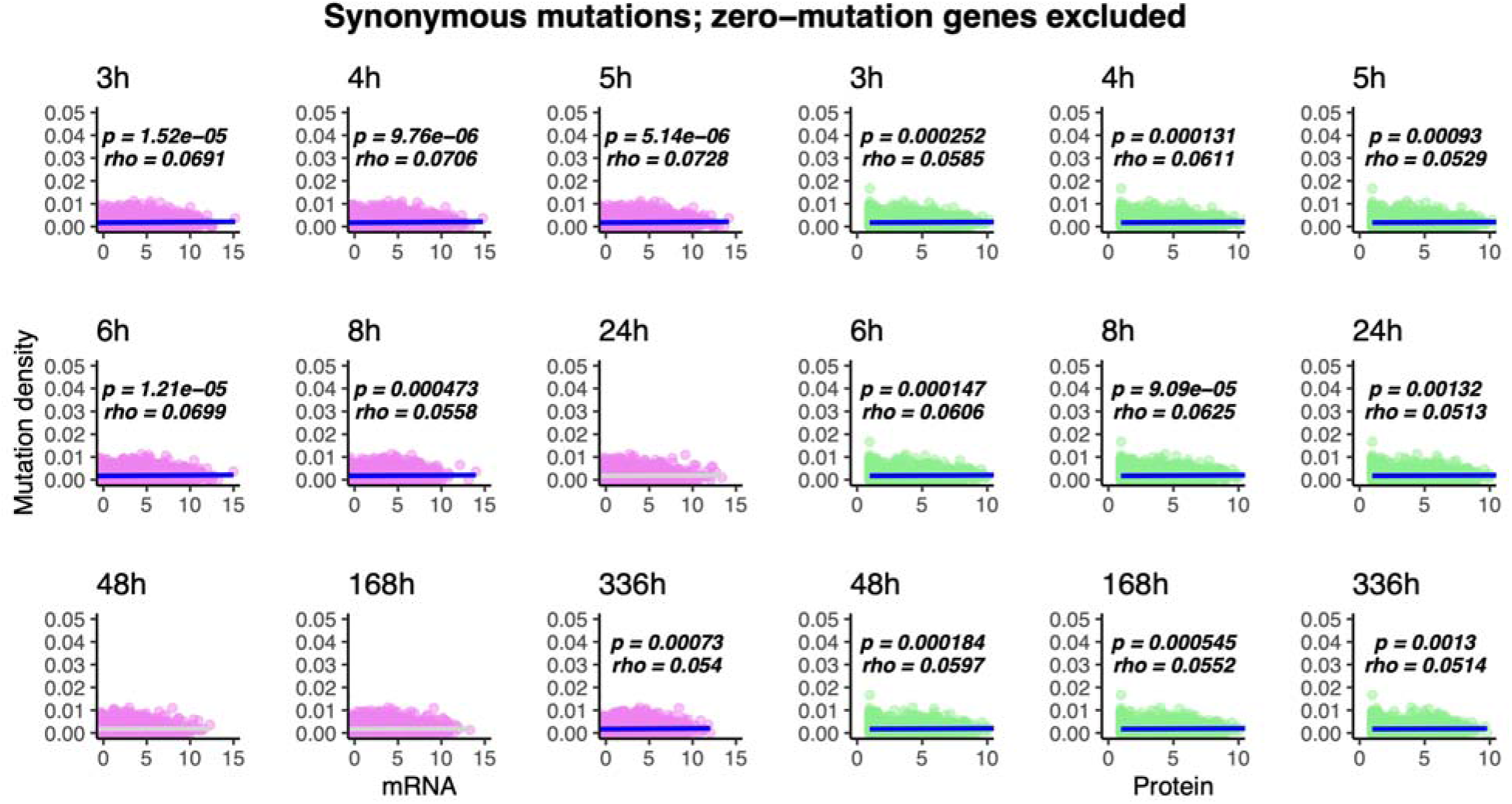
The density of observed synonymous mutations per gene across all hypermutator LTEE populations positively correlates with mRNA abundance in exponential growth phase, and positively correlates with protein abundance at all time points, when genes with no mutations are excluded. See legend to Supplementary Figure S1 for further details.

**Supplementary Figure S4.**
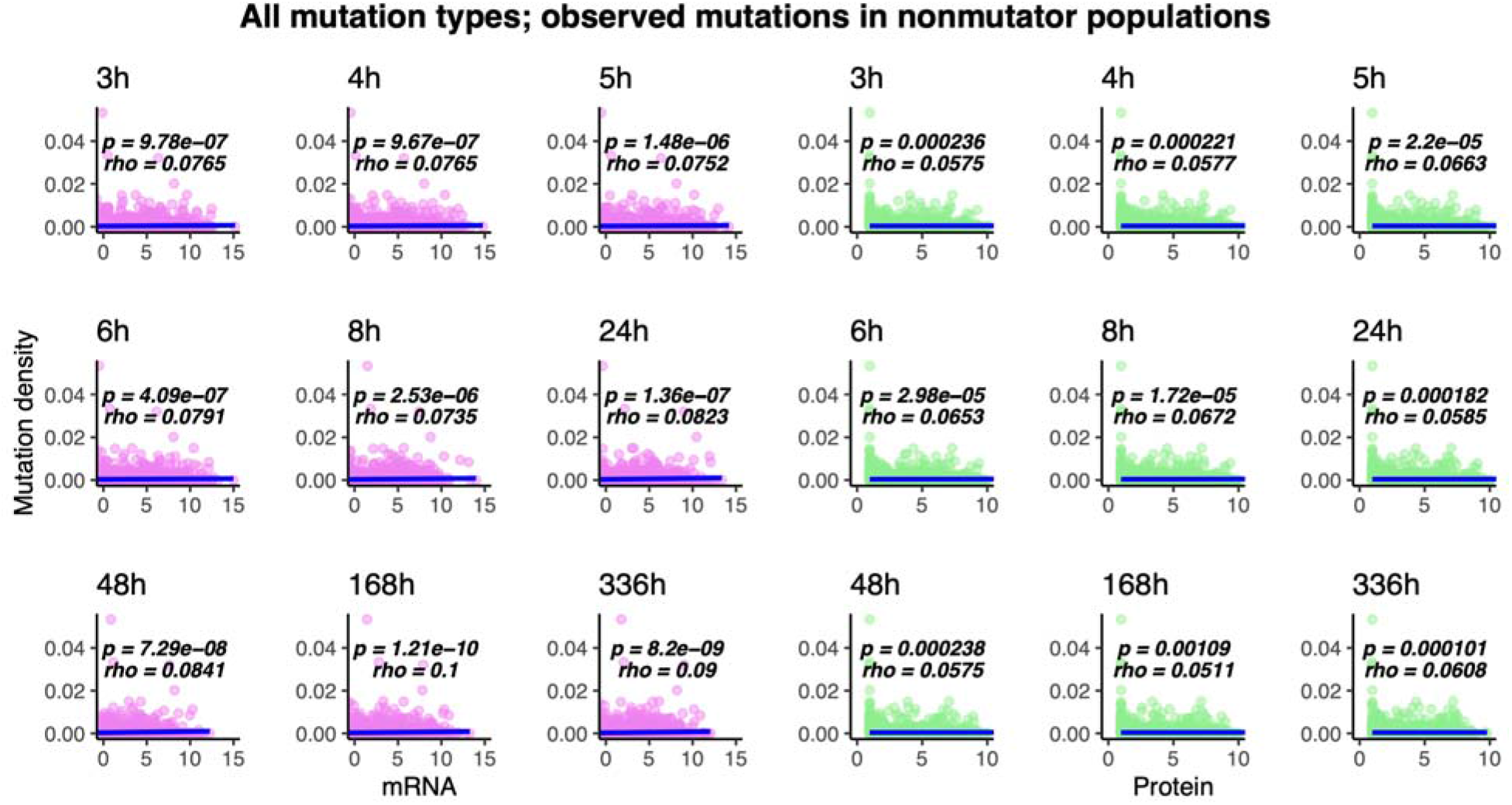
The density of observed mutations per gene across all nonmutator LTEE populations positively correlates with mRNA and protein abundance at all time points. See legend to Supplementary Figure S1 for further details.

**Supplementary Figure S5.**
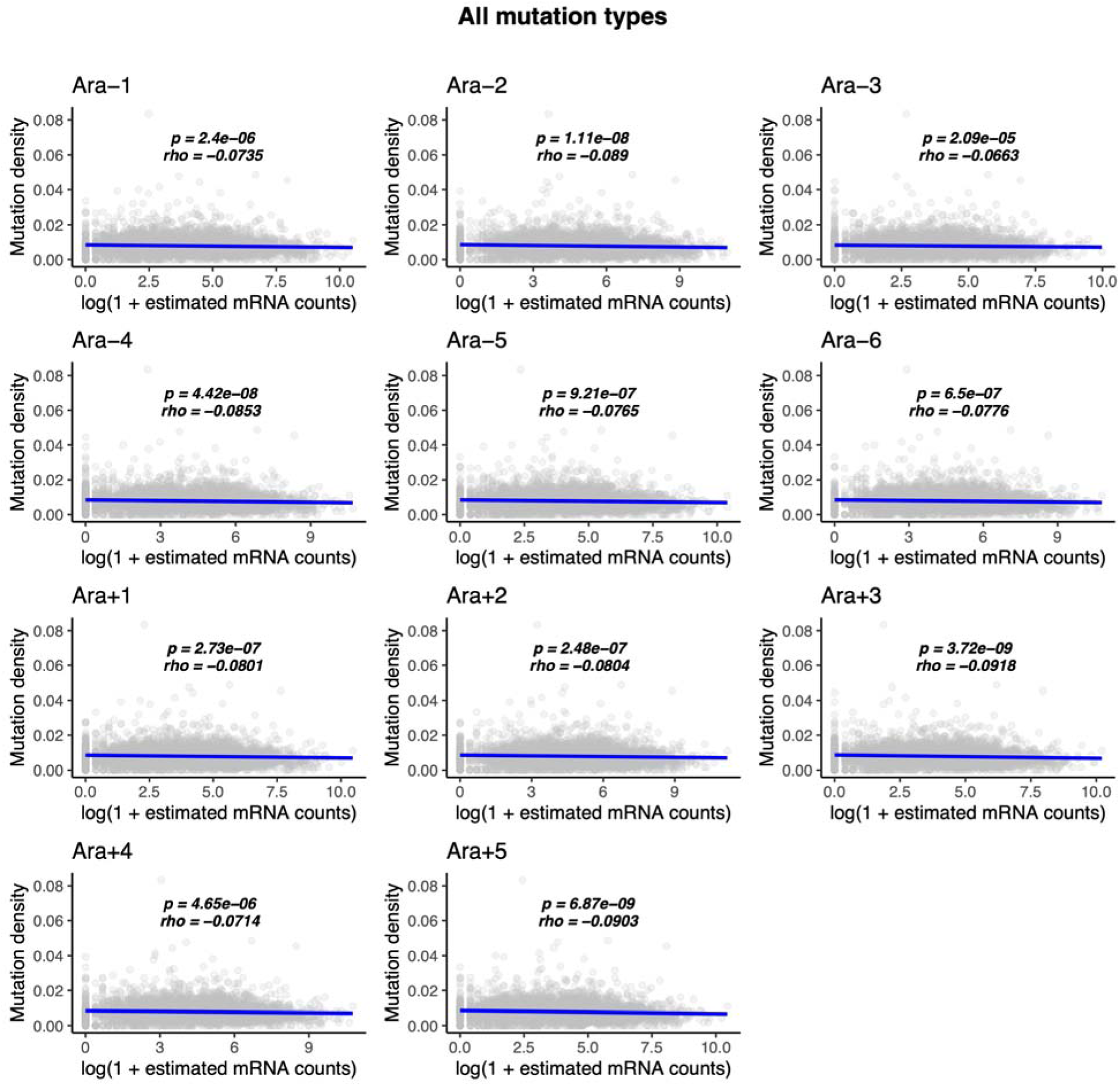
The density of observed mutations per gene across all hypermutator LTEE populations at 60,000 generations anti-correlates with mRNA abundance in exponential phase for all 11 50,000 generation LTEE clones grown in DM4000 media. These transcriptomic data were reported by Favate et al. (2021). Statistically significant correlations are shown in blue, while non-significant correlations are shown in light gray. Spearman correlation coefficients (*rho*) and associated *p*-values are shown on each panel.

**Supplementary Figure S6.**
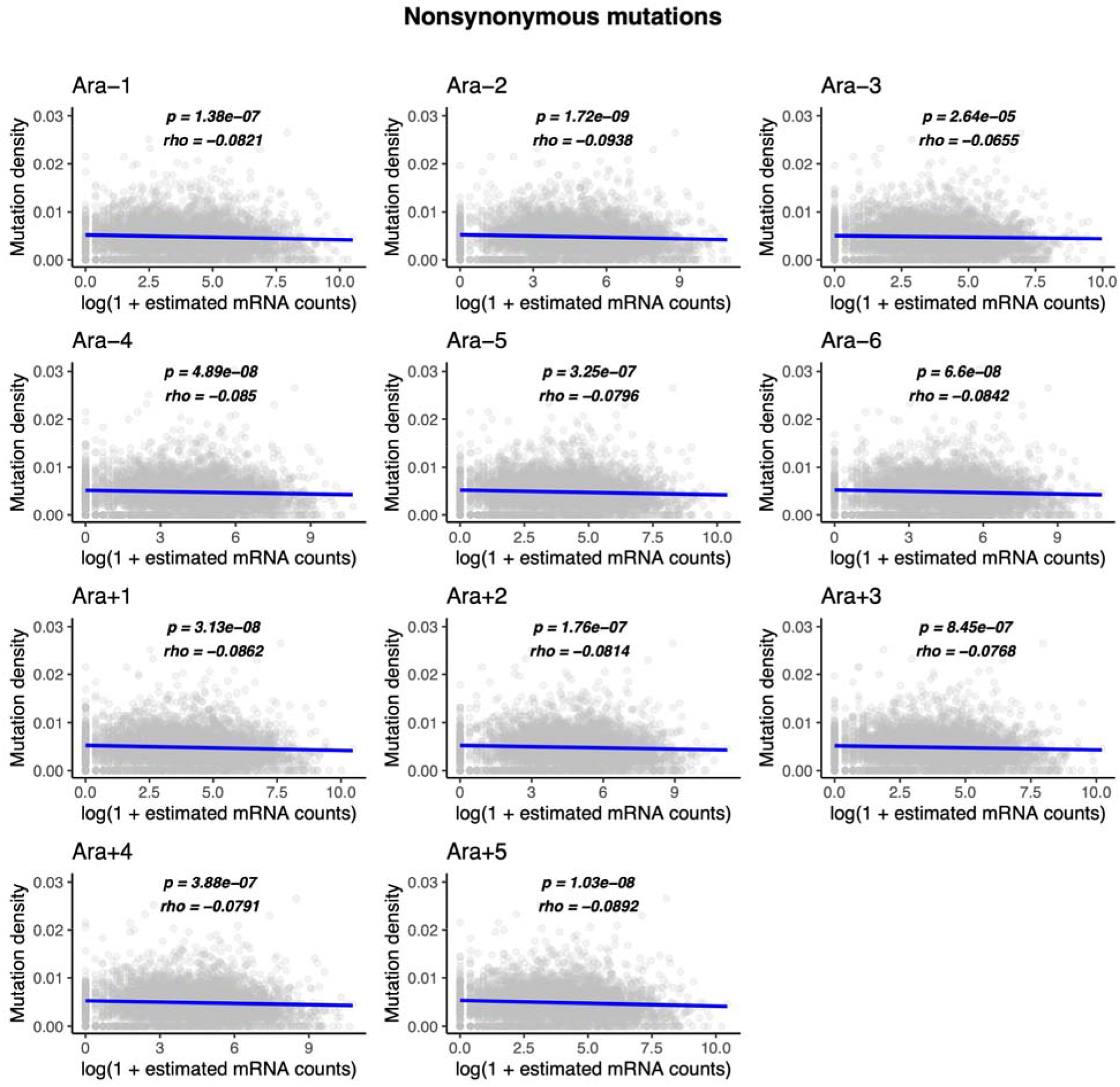
The density of nonsynonymous mutations per gene across all hypermutator LTEE populations at 60,000 generations anti-correlates with mRNA abundance in exponential phase for all 11 50,000 generation LTEE clones grown in DM4000 media. These transcriptomic data were reported by Favate et al. (2021). See legend to Supplementary Figure S5 for more details.

**Supplementary Figure S7.**
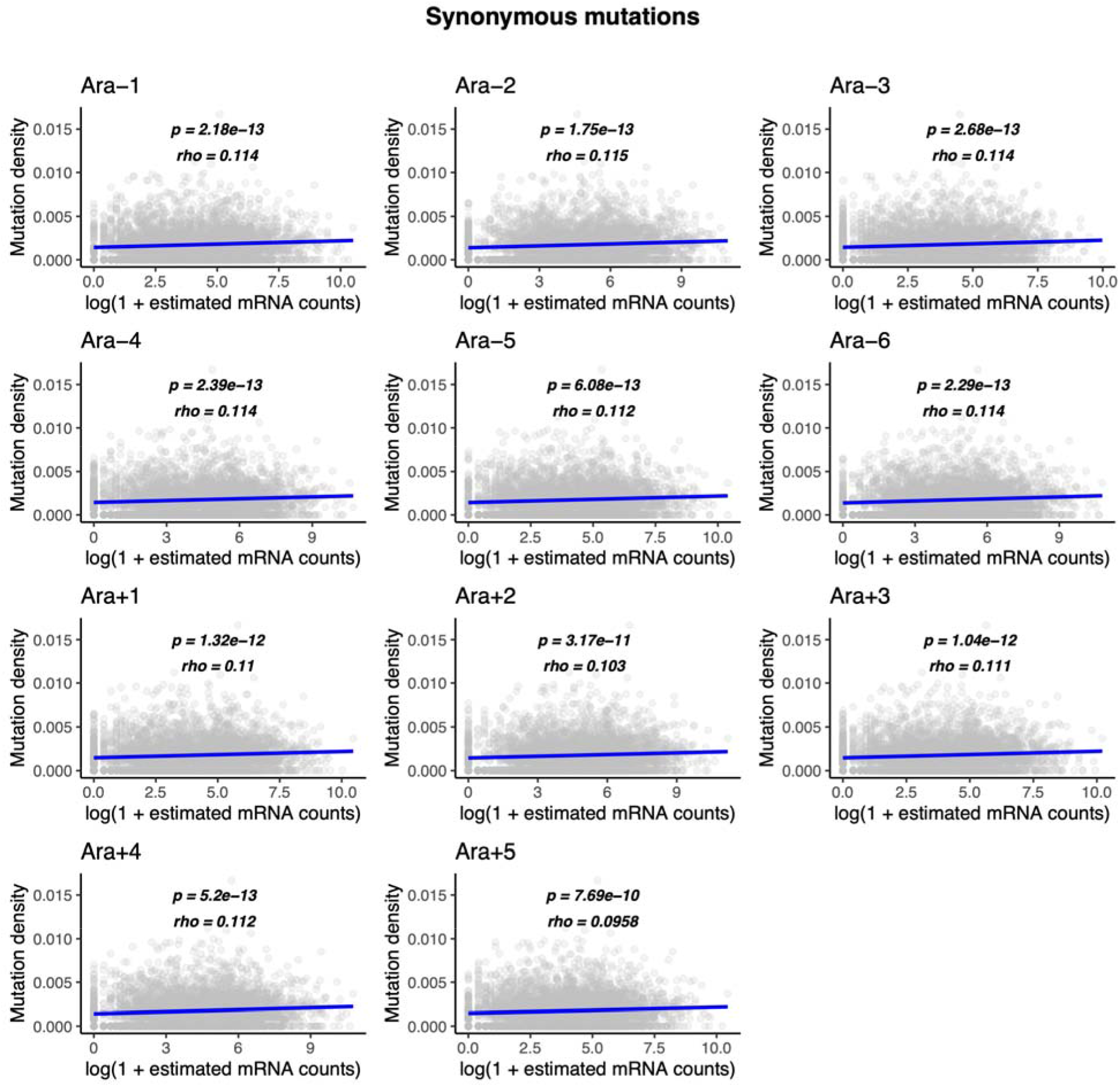
The density of synonymous mutations per gene across all hypermutator LTEE populations at 60,000 generations positively correlates with mRNA abundance in exponential phase for all 11 50,000 generation LTEE clones grown in DM4000 media. These transcriptomic data were reported by Favate et al. (2021). See legend to Supplementary Figure S5 for more details.

**Supplementary Figure S8.**
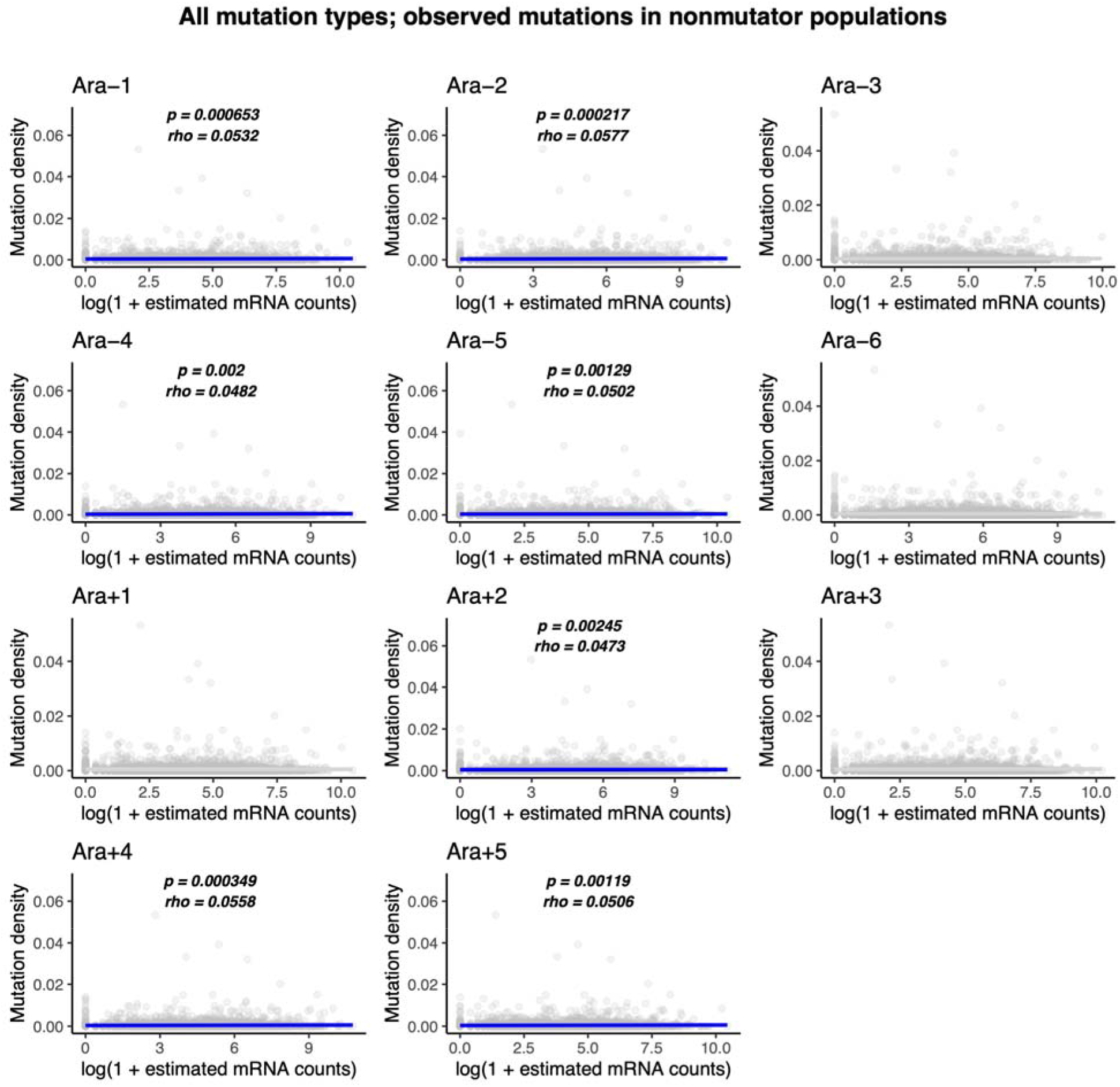
The density of observed mutations per gene across all nonmutator LTEE populations at 60,000 generations positively correlates with mRNA abundance in exponential phase in most of the 11 50,000 generation LTEE clones grown in DM4000 media. These transcriptomic data were reported by Favate et al. (2021). See legend to Supplementary Figure S5 for more details.

**Supplementary Figure S9.**
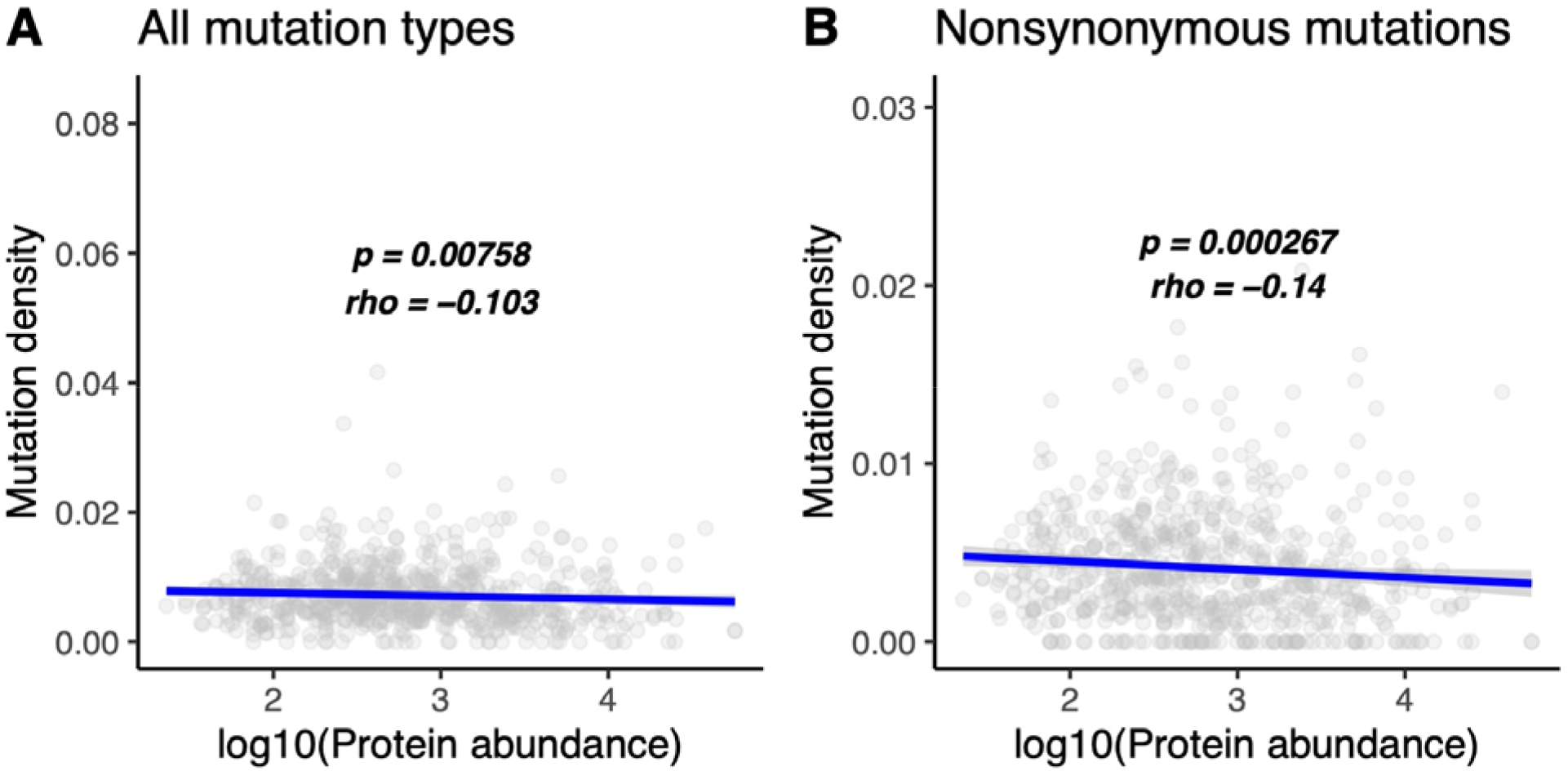
The density of observed mutations per gene across all hypermutator LTEE populations at 60,000 generations anti-correlates with protein abundance data in the ProteomeVis database. The ProteomeVis database is described in Razban et al. (2018). Each point corresponds to one of 664 genes with abundance data in ProteomeViz. At the time of accession, ProteomeVis did not contain abundance data for the remaining 3541 genes in the 60,000 generations LTEE metagenomics data (Good et al. 2017). Corresponding results for synonymous mutations in the hypermutator LTEE populations, and for all mutation types in the nonmutator LTEE populations, are not statistically significant.

**Supplementary Figure S10.**
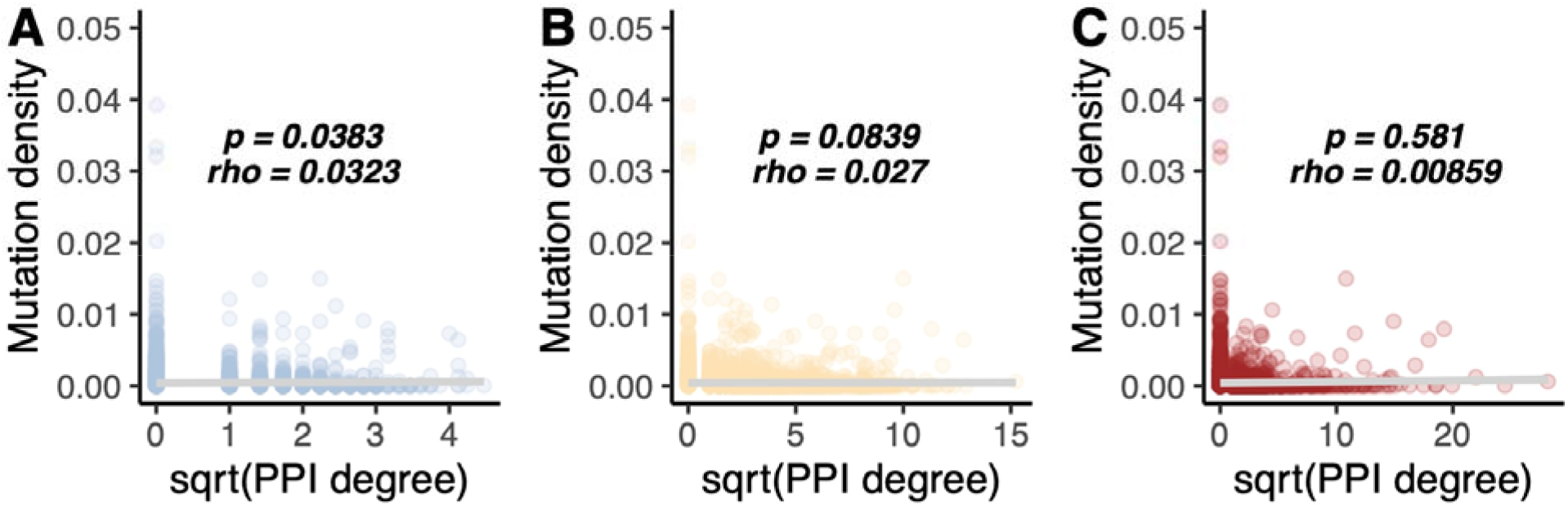
The density of observed mutations per gene across all nonmutator LTEE populations does not correlate with PPI degree. A) Comparisons to the PPI data from Cong et al. (2019) are shown in light blue B) Comparisons to the PPI data from Zitnik et al. (2019) are shown in orange. C) Comparisons to the ProteomeVis PPI dataset are shown in red.

